# An initial assessment of the sustainability of waterbird harvest in the United Kingdom

**DOI:** 10.1101/2022.03.02.482631

**Authors:** Matthew B. Ellis, Tom C. Cameron

## Abstract

1. There is a need to assess the sustainability of wild bird harvest in the United
Kingdom (UK), and more widely, across Europe. Yet data on populations and harvest sizes are limited.
2. We used a Demographic Invariant Method (DIM) to estimate Potential Excess Growth (PEG) for populations of UK wintering waterbirds and calculated a Sustainable Harvest Index (SHI) for each. We compared this with population trends and conservation classifications (e.g. Birds of Conservation Concern; BoCC) to assess the sustainability of harvests and the utility of these classifications.
3. Our approach found evidence for potential overharvest of mallard *Anas platyrhynchos*, Eurasian teal *Anas crecca*, gadwall *Mareca strepera*, Canada geese *Branta canadensis*, greylag geese *Anser anser* and woodcock *Scolopax rusticola*. Whether DIM methods predict overharvest is highly dependent on estimates of maximum population growth rates inferring PEG. We found estimates of maximum population growth to be variable across a range of different methods.
4. We found no relationship between SHI and short-term wintering trends or conservation classification under the UK’s BoCC framework. There was however a positive relationship between SHI and long-term wintering trends. *Policy Implications*: Our results suggest that UK based harvest is unlikely to be a major determinant of population trends for the majority of UK overwintering waterbirds, but harvest rates for some species may exceed that required to maintain stationary population growth. The lack of a relationship between conservation classifications and SHI strongly suggests that such conservation classifications are not an appropriate tool for making decisions about harvest management. Instead, our assessment provides the basis for a framework to make evidence-based decisions on sustainable harvest levels in the face of incomplete data. There is currently no clear policy instrument in the UK to support such a framework via controls on either harvest effort or mortality of waterfowl.

## Introduction

Hunting of waterbirds in the UK with shotguns has occurred since at least the 1600s (Helme, 1614). Contemporary recreational hunting of waterbirds in the UK targets nine species of duck, four species of goose and three waders. Coot *Fulica atra* and moorhen *Gallinula chloropus* are also huntable but are rarely pursued and there are no modern estimates of the harvest of these species. The total annual harvest of ducks, geese and waders in the UK is estimated at approximately 1.4 million (Aebischer, 2019; PACEC, 2014). These same species, hunted in winter in the UK, breed across Northern, Eastern Europe and Russia and are hunted across their ranges and migration routes.

The British Association for Shooting and Conservation (BASC) is the largest representative body for shooting sports in the UK. BASC has approximately 8,000 wildfowling members, which is likely to represent the majority of self-identified wildfowlers hunting ducks and geese on UK coasts and estuaries. Of these wildfowlers approximately 25% hunt at least once per season, and they harvest an average of 8 birds each per season (Crown Estate, 2021; Ellis, 2014). This leads to an estimated coastal harvest of 16,000 waterbirds. The approximately 1.384 million remaining harvested waterbirds are harvested inland over a broad range of habitats, including artificially fed sites. Released mallard *Anas platyrhynchos* are a significant proportion of this inland harvest with current release estimates of approximately 2.6 million (Madden, 2021), and harvest estimates of approximately 950,000 birds per year (Aebischer, 2019). A significant proportion of other wild duck and goose species are also harvested in these inland areas due to the creation and provision of appropriate habitat, as well as supplementary feeding.

The rarity, localisation and population trends of birds in the UK has been regularly assessed since 1996 under the Birds of Conservation Concern (BoCC) methodology (Gibbons et al., 1996). Species are assigned the highest priority rating (Red>Amber) for any of the criteria they exceed such that, for example, a species which exceeded the Amber breeding range decline criterion and the Red breeding population decline criterion would be classified as Red-listed. Species which do not exceed any of the assessment criteria are placed on the Green list. A species with an increasing or stable UK population can be classed as Amber due to, for example, the UK holding an internationally important proportion of the European population– e.g. Pink-footed goose *Anser brachyrhynchus*. This approach is different than the better-known International Union for the Conservation of Nature (IUCN) Red List which only reviews the extinction risk for species (IUCN, 2020). This can lead to disparities due to differences in the scale of assessment. For example, the Eurasian woodcock *Scolopax rusticola* is classed as least conservation concern by the IUCN due to a very large and stable global population and large range but has Red status on the UK BoCC due to long and short term declines in UK breeding range.

Increasingly there have been calls to prohibit or restrict the hunting of species that are Red or Amber listed on BoCC (Thomas, 2021; Welsh Parliament, 2020). Similar calls drawing attention to the BoCC status of hunted birds have been made by social media influencers (Common, 2016; UK Government and Parliament, 2016). Currently all huntable ducks and geese in the UK, with the exception of tufted duck *Aythya fuligula* and Canada goose *Branta canadensis*, are BoCC Red or Amber listed, most commonly due to the UK holding >20% of the European wintering population, rather than due to declines in breeding or wintering populations. Apriori then, BoCC may not be the most appropriate metric to judge conservation risks from additional winter mortality, but this has not yet been tested. Likewise, there have been calls to prohibit or restrict harvest in nationally and internationally protected sites. Typically, on these sites, harvest data is robust, and harvests are low (approximately 1% of the estimated UK harvest), compared to unprotected inland sites where regulatory authorities currently have limited room to act. These regulatory disparities lead to conflicts and this calls for an assessment of the sustainability of the UK harvest of such highly mobile species, that are constantly responding to food availability, recreational disturbance and hunting pressure with shifting range size and range location, at an appropriate spatial scale (Beatty et al., 2014; O’Connell et al., 2007).

In the absence of regular and systematic monitoring of geographic population structures or harvest returns for huntable species in the UK it has historically been difficult to assess the sustainability of current harvest levels, particularly so for data poor ducks compared to more regularly monitored European goose populations (Holopainen et al., 2018; Madsen et al., 2015). However, the development of methods allowing for estimates of harvest levels in the UK (Aebischer, 2019), and the development of analytical tools to assess the sustainability of harvest with incomplete demographic data now allows for initial assessments to be made even in the face of uncertainty (Eraud et al., 2021). The UK is party to the Agreement on the Conservation of African-Eurasian Migratory Waterbirds (AEWA) which is developing mechanisms to ensure flyway-level harvest sustainability and in the long term will be a more appropriate venue to make these decisions (Madsen et al. 2015). In the absence of these mechanisms we present an initial assessment of the sustainability of winter hunting mortality of UK waterbirds using best-available data, whilst recognising the likely sources of error and identifying areas where further research would be particularly valuable.

## Materials and Methods

We collated mean and confidence intervals for harvest estimates, population size estimates, and demographic data, and also population trends on huntable ducks, geese and waders in the UK. This included nine species of ducks (mallard *Anas platyrhynchos*; Eurasian teal *Anas crecca*; Eurasian wigeon *Mareca penelope*; gadwall *Mareca strepera*; northern pintail *Anas acuta*; northern shoveler *Spatula clyptea*; tufted duck *Aythya fuligula;* common pochard *Aythya ferina;* and common goldeneye *Bucephala clangula*), three waders (Eurasian woodcock *Scolopax rusticola;* common snipe *Gallinago gallinago*; and golden plover *Pluvialis apricaria*) and three goose species comprising a total of four goose populations (British (resident) and Icelandic (migratory) greylag goose *Anser anser*; Greenland/Icelandic pink-footed goose *Anser brachyrhynchus*; and resident Canada goose *Branta canadensis*). These data were used to estimate a Sustainable Hunting Index using a stochastic simulation approach using the “popharvest” R package (Eraud et al. 2021).

### Estimating harvest sustainability

A Sustainable Hunting Index (SHI + 95% confidence interval (CI)) was calculated for each species using the potential excess growth (PEG) function in the popharvest R package (Eraud et al., 2021). This index is calculated as the ratio of harvest to PEG such that any value in excess of 1 indicates overharvest and values under 1 are increasingly more likely to be sustainable as they approach zero. This is not unlike a surplus production model approach that is commonly applied to data poor or initial assessments for harvest of other vertebrate systems such as fisheries (Jennings et al., 2008).

The PEG for each species was calculated using a Demographic Invariant Method (DIM) which allows estimation of maximal potential population growth rates (λ_max_) with relatively few demographic parameters including age at first reproduction, life-history strategy categorisation and adult survival (Ernaud et al., 2021; Johnson et al., 2012). This stochastic simulation approach allows us to best account for the uncertainty in data sources.

The choice of life history strategy (“long” versus “short”) and the use of adult survival estimates from either ringing studies or estimated from species mass can have significant effects on the estimate of SHI. Given the uncertainty around these variables and the significant effects each can have on the estimated outputs we ran four models per species, varying life history strategy (“long” versus “short”) and adult survival (“reported” versus “estimated”) whilst holding other variables the same and present the model averaged results. In each model we ran 10,000 Monte Carol simulations per species with both population and harvest estimates drawn from a uniform range equal to the 95% confidence intervals of published estimates (full details on data sources given in supplementary online materials). The age at first reproduction is poorly understood for most species so was taken to be fixed at 1 for all duck species, 2 for all waders and 3 for all geese (Robinson, 2005). Distributions for drawing simulation parameters are given in the supplementary online materials and were guided by those recommended by the package authors (Eraud et al., 2021).

Models using “reported” adult survival use the average adult survival reported from ring returns. However, this can underestimate maximal survival rates under optimal conditions for harvested species and bias low the estimated sustainable harvest index. In models with “estimated” adult survival we provided body mass information to the model and allow it to draw on an estimate of adult survival in optimum conditions (Johnson 2012) from estimates of how adult survival responds to variation in body mass across many bird taxa (Ricklefs, 2000).

The popharvest package includes a safety factor (F_s_) which can be set from 0-1 to limit harvest to only a proportion of the PEG (with a recommended maximum of F_s_ = 0.5). Based on published recommendations (Dillingham & Fletcher, 2008), we set F_s_ = 0.5 for all species listed as least concern by IUCN. Pochard is listed as Vulnerable under IUCN and therefore we assigned an F_s_ = 0.1. Setting F_s_ to a recommended maximum of 0.5 does not well represent the sustainability of a harvest where hunters are being encouraged to maximise their harvest, for example on non-native geese or geese with expanding populations, e.g. Canada or pink-footed goose. We shall examine this issue later in the discussion. As well as calculating a confidence interval for SHI the package also calculates the probability that the current level of harvest is unsustainable. Estimated λ_max_ using published survival estimates from ringing studies and the inbuilt body mass predicted adult survival were compared with other published assessments for Canada goose and mallard to make sure our DIM approaches were producing reasonable estimates (Table S1).

We tested for relationships between SHI and Birds of Conservation Concern category using a Welch’s one-way ANOVA which allows for unequal variance and between SHI and wintering population trend (separately for short and long term) using linear regression. All analyses were conducted in R version 4.1.2 (R Core Team, 2021).

## Results

The estimated potential excess growth (PEG), sustainable hunt index (SHI) and probability of unsustainable harvest averaged across the four models can be found in Table 1 (individual model estimates presented in Tables S2-S5). The confidence intervals of the SHI estimates for the current harvest for mallard, teal, gadwall, Canada goose, both British and Icelandic greylag goose and both resident and migratory woodcock included values greater than 1 and so were predicted to be potentially unsustainable against an objective of harvesting 50% or less of maximum annual productivity. Harvest levels for some species, e.g. Canada goose, were up to eight times greater than half the potential excess growth (i.e.when F_s_ is set to 0.5). The highest predicted SHI for duck species was for Eurasian teal at 1.069 (95% CI 0.423 – 2.088), followed by mallard, gadwall, shoveler and wigeon in decreasing order (Table 1).

**Table 1:** The estimated mean λ_max_ (calculated as exp(R_max_)), potential excess growth (PEG; reported to two significant figures), sustainable hunt index (SHI) and the probability of current harvest levels being unsustainable with a safety factor (F_s_) = 0.5 for huntable waterbird species populations overwintering in the United Kingdom, except for pochard where F_s_ = 0.1. The presented figures are an average of four models using either a “long” or “short” life history strategy and either the “reported” or “estimated” adult survival rates.

| Species | Estimated $\lambda_{\max}$<br>(95% CI) | Mean Potential Excess<br>Growth (95% CI) | Mean Sustainable<br>Hunt Index (95% CI) | Probability of<br>unsustainable harvest |
| --- | --- | --- | --- | --- |
| Mallard | 1.871<br>(1.435 - 2.826) | 1,000,000<br>(590,000 – 1,700,000) | 1.057<br>(0.471 - 1.950) | 0.504 |
| Teal | 2.041<br>(1.515 - 3.469) | 160,000<br>(87,000 – 270,000) | 1.069<br>(0.423 - 2.088) | 0.512 |
| Wigeon | 1.993<br>(1.461 - 3.36) | 160,000<br>(78,000 – 280,000) | 0.414<br>(0.111 - 0.971) | 0.018 |
| Gadwall | 1.774<br>(1.442 - 2.922) | 8900<br>(5700 – 17,000) | 0.772<br>(0.240 - 1.620) | 0.248 |
| Pintail | 1.840<br>(1.454 - 3.067) | 6100<br>(3400 – 11,000) | 0.221<br>(0.038 - 0.538) | 0.000 |
| Shoveler | 1.949<br>(1.470 - 3.158) | 6500<br>(3500 – 12,000) | 0.426<br>(0.117 - 0.981) | 0.020 |
| Tufted duck | 1.798<br>(1.458 - 2.452) | 41,000<br>(26,000 – 63,000) | 0.155<br>(0.056 - 0.317) | 0.000 |
| Pochard | 1.852<br>(1.442 - 2.715) | 1800<br>(980 - 3000) | 0.359<br>(0.082 - 0.859) | 0.001 |
| Goldeneye | 1.713<br>(1.449 - 2.450) | 5600<br>(3700 - 9500) | 0.201<br>(0.026 - 0.474) | 0.000 |
| Canada goose | 1.223<br>(1.164 - 1.301) | 17,000<br>(12000 - 22000) | 2.148<br>(0.804 - 3.997) | 0.926 |
| Greylag goose<br>(Icelandic) | 1.199<br>(1.158 - 1.281) | 8300<br>(6700 – 11,000) | 2.777<br>(1.937 - 3.541) | 1.000 |
| Greylag goose<br>(British) | 1.200<br>(1.158 - 1.282) | 13,000<br>(10,000 – 17,000) | 4.163<br>(0.869 - 8.123) | 0.956 |
| Pink-footed<br>goose | 1.205<br>(1.175 - 1.253) | 47,000<br>(35,000 – 62,000) | 0.448<br>(0.072 - 0.906) | 0.006 |
| Woodcock<br>(Resident) | 1.411<br>(1.302 - 1.61) | 28,000<br>(16,000 – 46,000) | 0.711<br>(0.394 - 1.183) | 0.097 |
| Woodcock<br>(Migratory) | 1.411<br>(1.302 - 1.61) | 210,000<br>(120,000 – 340,000) | 0.636<br>(0.353 - 1.057) | 0.042 |
| Common Snipe | 1.467<br>(1.336 - 1.732) | 210,000<br>(140,000 – 320,000) | 0.392<br>(0.133 - 0.763) | 0.000 |
| Golden plover | 1.841<br>(1.545 - 2.564) | 130,000<br>(81,000 – 200,000) | 0.018<br>(0.002 - 0.043) | 0.000 |

Using the mean and CI of SHI estimates we can also estimate how probable it is that a current harvest is unsustainable. For wintering UK ducks, that is harvest greater than 50% of the maximum potential surplus production of current winter population sizes, probabilities were generally low, even for mallard (0.504) and teal (0.512). For pochard, where the threshold for sustainability was set to only 10% of the maximum surplus growth due to recent breeding population declines in continental Europe as well as poor UK breeding and harvest data, SHI was 0.359 (95% CI 0.082 – 0.859) with a low probability of unsustainable harvest (0.001). Golden plover and common snipe did not have SHIs or 95% CIs exceeding 1 and the probability of an unsustainable harvest was low (Table 1). However, the confidence intervals for both woodcock populations included values in excess of 1, though with low probabilities of unsustainable harvest (0.097 and 0.042 for resident and migratory populations respectively).

The choice of life history strategy (e.g. short vs long) and survival estimates (e.g. reported vs DIM estimate) had a large effect on the magnitude of model-specific SHI estimates, but had little effect on the overall species order (Figure 1 and Tables S2-S5).

**Figure 1:**
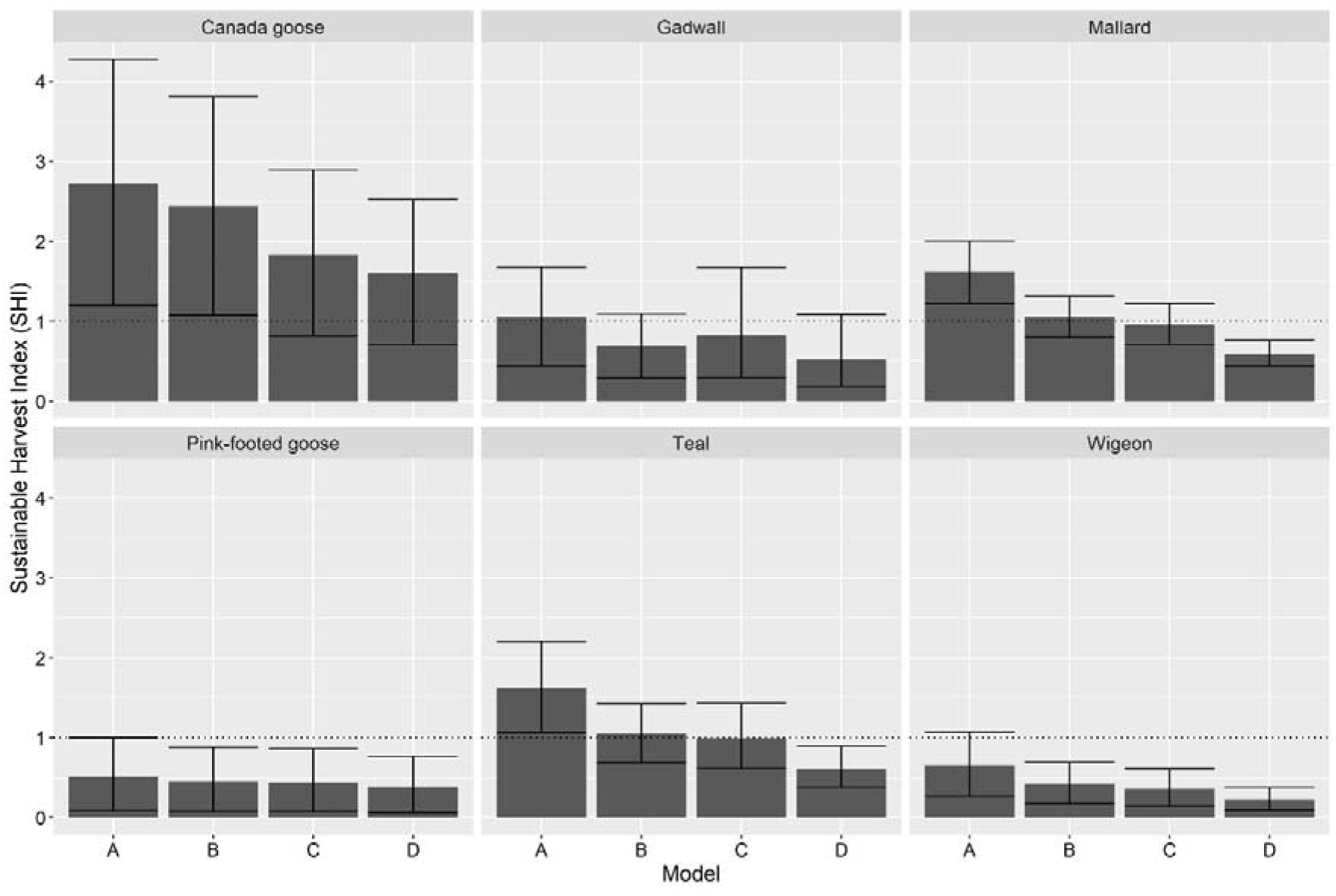
Mean (and 95% confidence interval) sustainable harvest index (SHI) for selected huntable waterbirds in the United Kingdom modelled using survival rates estimated from mass and (A) a long life history or (B) a short life history, and survival rates reported in the literature and a (C) long life history or (D) short life history. SHI is calculated with F_s_ = 0.5 for all species and the dotted line at SHI=1 represents the threshold for overharvest. Species were selected to represent the most well studied huntable species.

There was no significant relationship between the average SHI and the BoCC5 category across all species (F(_2,4.6636_) = 1.2374, p = 0.371), and there was also no significant relationship between average SHI and short-term wintering trends (β = 0.007, p = 0.446). There was a positive relationship between average SHI and long-term wintering trends (β = 0.008, p = 0.040)(Figure 2) such that the species with the greatest population growth had the highest predicted average SHI.

**Figure 2:**
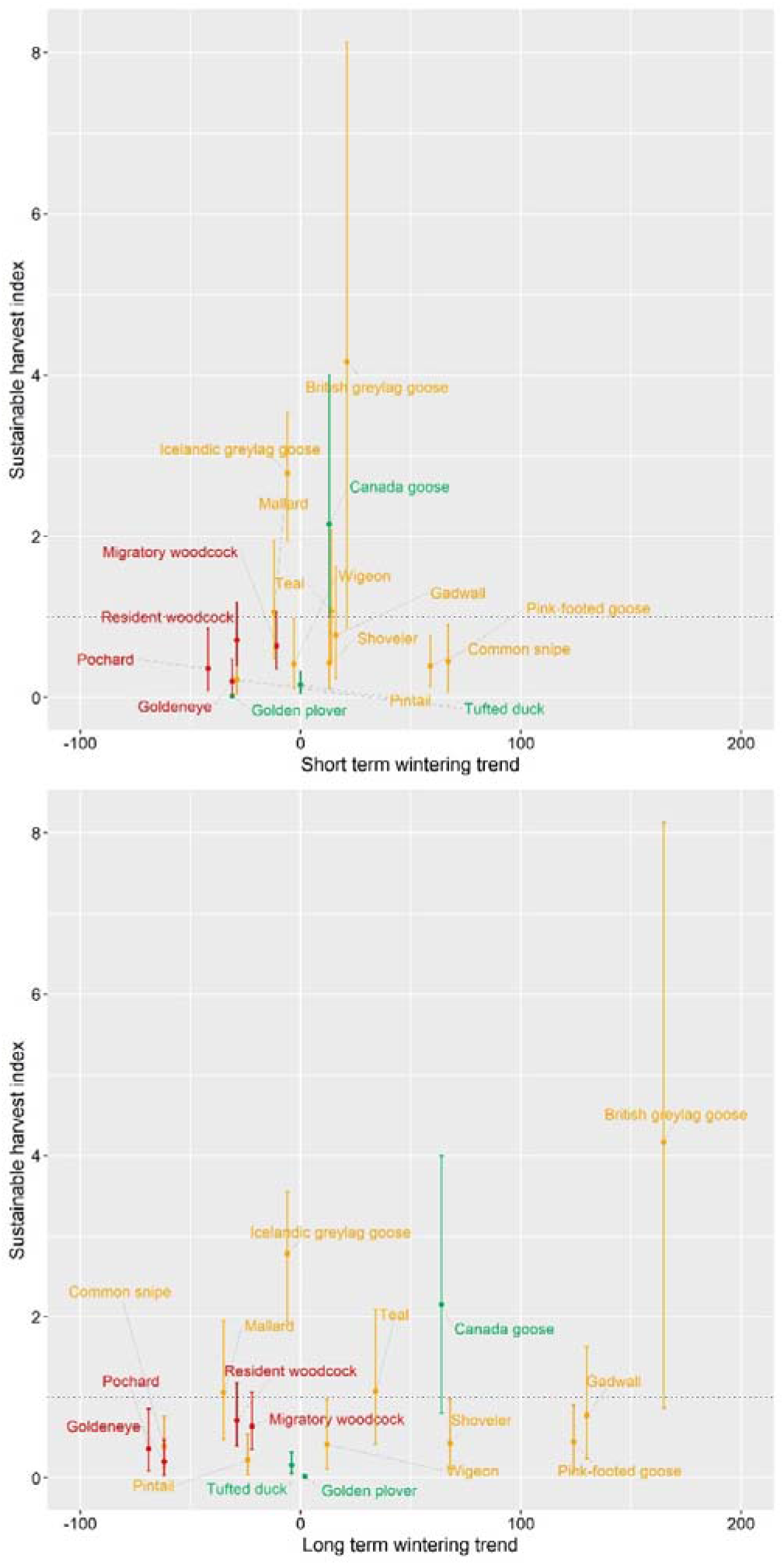
Mean and 95% confidence intervals for the sustainable harvest index (SHI) for huntable waterbirds in the United Kingdom along with their short-term (2008-2018) and long-term (1970 – 2018) wintering population trends. The dotted line at SHI = 1 represents the threshold for overharvest such that species falling above the line are likely to be experiencing overharvest. SHI is calculated with F_s_ = 0.5 for all species other than pochard where F_s_ = 0.1. Species are colour coded according to their classification under Birds of Conservation Concern 5.

## Discussion

In this study we used data on estimated waterbird population sizes and harvest levels to calculate sustainable harvest indices (SHI). For the majority of ducks and waders hunted in the UK we found no evidence of unsustainable harvests, but we recommend further scrutiny for mallard, teal, gadwall and woodcock where the SHI confidence intervals included values in excess of 1. For this assessment we used a moderating factor (F_s_) of 0.5 for all ducks and waders other than pochard. If a socio-ecological management plan for duck or wader harvest in the UK and Europe aimed for higher or lower harvests than 50% of the surplus production, then the SHI assessment would change.

Our estimates of SHI for wild geese suggest that Canada geese and Icelandic and British greylag geese may be subject to unsustainable harvest levels in the UK. This remains the case for Canada goose and both greylag goose populations if a higher moderating factor is chosen (F_s_=0.9 where harvest target is 90% of maximum surplus growth). However, British greylag goose, Canada goose and indeed gadwall experience strongly increasing population trends in recent years (Frost et al., 2019). Although there is some evidence of levelling off of these trend increases in the most recent years (Frost et al., 2021), particularly for Canada goose which could indicate that the curent harvest is finally having an effect on stabilising or reducing its numbers. In this context an unsustainable harvest of non-native Canada goose may be desirable but would suggest management to reduce that harvest may be required at some point.

Our SHI estimate for the migratory Icelandic greylag goose population is high with a high probability of unsustainable harvest. The Icelandic greylag goose population trend is declining, and this species is subject to significant additional hunting mortality outside of the UK. However, there are difficulties in accurately assessing the proportion of the harvest to assign to the resident and migratory populations and between the two main nations that hunt this population (UK and Iceland), which can clearly have a significant impact on the calculation of the SHI. For example, the mean annual harvest of greylag geese in Iceland was 43,000 per annum from 2015/16 to 2019/20 (Statistics Iceland, 2022), and we estimated the harvest for the UK as 22,500, giving a total estimated harvest across the flyway of 65,500 from a total estimated pre-breeding population of 60,000 (Brides et al., 2021). This suggests that action to improve estimates and limit the total harvest of this population would be warranted, at least as an adaptive experiment to test for linkage between harvest reduction and population recovery.

The high SHIs and 0.5 probabilities of unsustainable harvest for mallard and teal indicate that these species may be subject to unsustainably high levels of hunting mortality and this appears to accord with slowing of growth rates in teal populations and ongoing declines in mallard (SHI 95% confidence intervals included estimates greater than 1 in three out of the four models we ran for both species; Tables S2-S5). For teal we may be underestimating the population size from which UK hunters’ harvest, this is due to their high rate of turnover (such as seen in France; Pradel et al., 1997) and a generally more “open” population than we have traditionally considered (Calenge et al., 2010). Our apriori best candidate model for mallard and teal uses a short life history and DIM estimates of adult survival (Table S5), with maximum adult survival values that are reasonable from our review of the literature (Table S1) also predicts mean SHI values above 1. This result highlights an immediate need to improve population demographic data and reduce harvest, as harvest reduction can help us understand how populations respond to harvest control and therefore better inform and manage the sustainability of future hunting opportunities, as, for example, seen with mallard in North America (Smith and Reynolds, 1992).

We found no relationship between SHI and short-term wintering population trends and a positive trend with SHI and long-term wintering trends such that those species currently predicted to exposed to the most unsustainable levels of take were demonstrating the greatest population growth. This latter result is caused by long term high growth rates in goose populations now experiencing high harvest mortality. We suggest that this indicates that hunting of waterbirds in the UK is not a primary driver of population trends. This is to be expected as waterfowl harvests tend to be a function of waterfowl productivity in a given year (Fox et al., 2016; Holopainen, Christensen, et al., 2018). This means hunters tend to harvest more waterbirds in years when surplus production is higher and moderate harvests tend not to have long term effects on waterfowl population sizes. That is not to say that the sum of harvests across the flyway would give similar results, and we encourage further work using these tools but on international scales. Likewise further scrutiny of trends and over what period they apply may be useful as long-term trends for the total Eurasian woodcock population size (e.g. 25yr), were stated as stable in 2015 but are now stated as having strong declines (Birdlife 2015 vs 2021). Immediate steps would be to build further realism into this initial model assessment of sustainability by incorporating the demographic breakdown of the hunting bag as affecting juvenile or adult survival and attempting the model with estimates of population sizes and hunting mortality at larger spatial scales across the UK, mainland Europe and western Russia.

The lack of a relationship between SHI and BoCC5 categories suggests that classification as Red or Amber listed by this scheme is not an appropriate tool to use as an aid to harvest management decisions. This is not unexpected as BoCC5 classification depends on a variety of factors including breeding and wintering localisation and is not solely dependent on international population trends or demographic sensitivities of the species concerned. For example, pink-footed geese are amber listed for non-breeding localisation (greater than 50% of the UK non-breeding population at ten or fewer sites) and non-breeding international importance (UK holds at least 20% of the European non-breeding population)(Stanbury et al., 2021), despite significant long and short term population increases (Burns et al., 2020). However, this does not undermine the value of BoCC as a useful tool for the initial prioritisation of species of potential conservation concern at appropriate spatial scales; particularly for making habitat availability and restoration decisions.

In estimating the potential excess growth or annual recruitment in a bird population with which to assess the sustainability of harvest, a safety feature has been included so that the threshold with which to judge overharvesting is a proportion of the excess growth (Eraud et al., 2021; Johnson et al., 2012; Niel & Lebreton, 2005). This is wise due to well-recognised risks in using surplus growth approaches in data poor systems to maximise harvests (Jennings et al., 2008). The sequential use of this safety feature in frameworks designed for initial assessments of the sustainability of wild bird harvests has led to a clear and well described framework where the proportion of the excess growth that should not be exceeded by harvest can decline with data availability, population size, and declining conservation status (Dillingham & Fletcher, 2008; Eraud et al., 2021). But this initial assessment method does not take into account any specific harvest management objectives that may be equally important in utilitarian conservation systems such as legal recreational fishing or hunting, where the objective may be to maximise harvest opportunities for some species. This may be due to the species being non-native (e.g. Canada or Egyptian goose *Alopchen aegyptiaca* and Mandarin duck *Aix galericulata* in the UK), causing damage (e.g. pink-footed goose) or due to its relatively high abundance and productivity (e.g. Eurasian teal) making it a candidate for safe, sustainable regulated harvesting of wild food resources.

Harvesting up to 50% of the surplus production of highly productive species such as waterfowl is not unreasonable, where, in North America for example, 5.9 million Mallards were harvested from a breeding population of 14 million in 1999 (Wilkins & Cooch, 1999). Assuming the mean DIM estimated maximum population growth rate with a slow life-history of 1.765 for mallard, that represents 55% of the estimated surplus production in that year. Similar estimates of large harvest of 30+% of the annual surplus production of mallards in North America are common and sustained (e.g. U.S. Fish and Wildlife Service 2018). However, it is potentially informative to alter the level of precaution applied to the estimates of SHI. For example, reducing the F_s_ for Eurasian wigeon from 0.5 (reported in Table 1) to 0.1 changes the mean SHI estimate from 0.413 to 2.064. Likewise, these models can be used to estimate the harvest required to achieve an SHI value ≥1 for pink-footed goose if the aim is to reduce or stabilise population growth and expansion.

We have completed this initial assessment of the sustainability of waterbird harvest in the UK using the best available data and applying all potential estimates of maximum potential growth. However, we recognise that there are multiple sources of uncertainty in the data and parameter estimates. Population estimates of overwintering waterbirds in the UK are well documented and widely accepted, but there are known issues that influence reliability for this exercise. For example, the latest population estimates are scored for reliability out of 3 (3 = least reliable), and greylag geese, Canada geese, teal and goldeneye are scored 2 and mallard, snipe, woodcock and golden plover are scored 3 (Frost et al., 2019). Wetland bird survey (WebS) counts are made by an extensive network of volunteers overseen by the British Trust for Ornithology (BTO), and for many sites data extends back to 1947. However, coverage is incomplete and 39% of WeBS sites are currently not counted, of which 19.5% are “high” or “very high” priority (BTO, 2022). Such gaps undoubtedly contribute to the low reliability of some population estimates. Furthermore, changes to the methods used to make population estimates can have significant impacts. For example, the latest teal population estimates doubled from previous estimates as a result of new calculation methods (Musgrove et al. 2011; Méndez et al. 2015).

Overall, UK population estimates are likely to be underestimates due to count methods and the combined effects of hunting mortality, migration and turnover – this will result in underestimates of the potential excess growth and lead to a perceived greater proportion of the population being harvested. Depending on the management objectives (maximising population versus maximising sustainable harvest) this may be more or less important for decision makers.

Waterbird survival estimates are reported variously from the UK, North America or across Europe, but generally from ringing recaptures. The DIM approach requires good estimates of maximal adult survival under optimal conditions (Eraud et al., 2021). Underestimates of survival rates will bias estimates of population growth rates, potentially overestimating excess growth and hence underestimating the sustainable harvest index. We reviewed different approaches to estimate the maximum population growth for two of the most well studied hunted species to see if the model approaches produced “sensible” estimates (Table S1). We assumed Canada geese had a “long” life history and mallard “short” (equivalent to Table S2 and S3 respectively). The DIM estimate of maximum population growth using reported survival of 1.225 for Canada goose seems reasonable in light of studies across its range over varying time periods (1.07-1.74). However, for mallard, the DIM estimated maximum population growth using reported survival of 1.980 appears high relative to realised maximum population growth observations of 1.20-1.59 (Table S1). However, for both species taking published estimates of nesting attempts, nest success and brood survival can lead to productivity estimates more in line with the highest values e.g 1.58-2.10 for mallard (Table S1). As all realised population growth estimates from time series counts include hunting mortality, natural mortality and variation caused by environmental conditions and also suffer survey design error and biases, this may explain this discrepancy where observed trends are never estimating maximum population growth potential. Further work is required but given this review we feel the best estimates for Geese likely come from Table S2 and S4, and best SHI estimates for ducks and waders likely come from Table S5.

The accuracy of current UK waterbird harvest estimates is unknown (Aebischer, 2019) and developing a harvest recording system for the UK should remain a priority, not least to assist with management decisions at the flyway-scale (Madsen et al., 2015). The prospects for improving harvest data quality through voluntary systems seem low in the UK, especially in light of the diversity of hunters and hunter organisations, low trust in regulatory agencies by hunters and low levels of participation in other voluntary schemes (Ellis, 2020). However, it would seem to be of high policy importance to hunters and their organisations to be able to demonstrate the sustainability of their hunting activities. Under-reporting of hunter bags has been demonstrated to result in overestimates of national harvests and so positive engagement by hunters with bag recording schemes is more likely to result in positive outcomes for hunters than non-engagement (Aubry and Guillemain, 2019). Appropriate policy intervention may be welcome in this area provided it is applied to all participants, so that harvest data is collected from inland and private land areas as well as public coastal areas, areas of significantly higher and lower hunting pressure respectively.

Making decisions with incomplete data is difficult. We have used the best available data to make this preliminary assessment of the sustainability of harvest levels in the UK, but clearly further work is needed, including a structured assessment of the harvest at the flyway-scale. Our assessment provides a first step towards an informed discussion of harvest levels in the UK and whilst improved data collection will be useful, a collaborative approach to management based on the principles of Structured Decision Making and Adaptive Management could provide a transparent and trusted system to initiate with existing data that is “good enough” (Holopainen et al., 2018; Johnson et al., 2018; Williams & Brown, 2014). This approach is beginning to be developed in Europe by AEWA through mechanisms such as the European Goose Management Platform. This removes the need to rely on simplistic and inappropriate traffic light systems (such as Birds of Conservation Concern) which invariably result in poorly supported and simplistic outcomes (prohibiting versus status-quo) and provides a framework for countries like the UK with no specific harvest management policy instruments currently in place. At the same time we would recommend a move towards a system which explicitly values recreational hunting, such as how the UK values recreational fishing, and allows for an ecologically sustainable approach which is socially acceptable, evidence based and able to initiate with incomplete data.

## Authors’ Contributions

ME and TC conceived the idea, ME designed the methodology, collected and analysed the data, and wrote the first draft of the manuscript. TC contributed critically to drafts of the manuscript, validated the analyses and produced the supplementary information.

## Conflict of interest

MBE is employed by and is a member of BASC, the UK’s largest representative body for shooting sports. TCC is a member of BASC, the British Trust for Ornithology and Essex Wildlife Trust.

## Data availability statement

Data hosted on the DataDryad Digital Repository https://doi.org/10.5061/dryad.3tx95x6k2

## Notes

https://doi.org/10.5061/dryad.3tx95x6k2

### Competing Interest Statement

MBE is employed by and is a member of BASC, the UKs largest representative body for shooting sports. TCC is a member of BASC, the British Trust for Ornithology and Essex Wildlife Trust.

### Summary of Updates

Clearer reporting of methods and improved statistical methods

