## Supplemental material for "An initial assessment of the sustainability of waterbird harvest in the United Kingdom"

**Supplementary Online Material from Ellis & Cameron – an initial assessment of sustainability of waterbird harvest in the UK**

*Data sources*

Wintering bird populations for the UK were obtained from the latest published estimates (Frost et al., 2019). The estimate for mallard was increased by 2.6 million birds to account for the annual release of captive-raised birds for shooting purposes (Madden, 2021). A significant proportion of the UK mallard harvest is likley to be from these released birds, and a number of released birds will inevitably be included in the UK population estimates, but in the absence of further information on the breakdown of the mallard harvest we are unable to further refine our estimate. We estimated a standard deviation for the population size estimates by averaging the Wetland Bird Survey (WeBS; Frost et al. 2021) index for each species for the winters 2012/13 – 2016/17, which corresponded to the period used to estimate duck populations. We treated this mean index as equal to the estimate from Frost et al. (2019) and then calculated population estimates for each year based on their WeBS indices and calculated the standard deviation of these estimates. Population estimates for greylag goose, pink-footed goose, golden plover, snipe and woodcock were based on single years, but the same time frame was applied to standard deviation estimates in order to account for any interannual differences. Population estimates in the UK are based on five-year winter mean peaks (as opposed to mid-January counts used elsewhere in Europe) and are the counts used by the UK statutory authorities. We recognise that the population estimates are likely to be underestimates as they do not account for an unknown (and interannually variable) proportion of the birds either having been harvested or migrated prior to being counted. The latest (2016) harvest estimates for each population and their 95% confidence intervals were taken from Aebischer (2019).

Estimates of the population size and harvest of resident and migratory populations of greylag geese and woodcock are available separately and so we analysed these sub-populations independently. The resident woodcock population was estimated as three times the number of breeding males. This was subtracted from the total estimated overwintering woodcock population to provide an estimate of the migratory woodcock population. Resident woodcock harvest was estimated as equal to the proportion of breeding birds in the overwinter population (13%). The remaining 87% of total harvest was assigned to migratory woodcock. We recognise that further mortality of UK breeding woodcock will occur outside the UK in southern Europe, but we have no estimate of this mortality and can only consider the contribution of UK hunters to the harvest of birds that winter in the UK in this initial assessment. Population estimates for Icelandic and British greylag geese overwintering in the UK are reported separately and no adjustment is needed. An estimated harvest of migratory Icelandic greylag geese in the UK (Frederiksen, 2002) was deducted from the total UK greylag goose harvest to provide an estimate of the harvest of British greylag geese. However, it should be noted that the estimate was from 1996-2000 and no new estimates have been made.

Estimates of adult survival rates were generated from Robinson (2005) with reported standard errors multiplied by 1.96 to give an approximate 95% confidence interval. The average standard error for adult survival for all reported waterbirds (0.03) was used for species where no standard error was reported (e.g. wigeon, shoveler, greylag goose, woodcock and golden plover). We used the same survival estimates for both resident and migratory populations of woodcock and greylag geese. These adult survival estimates include mortality from hunting and so are likely to underestimate the maximum achievable survival rates under optimal conditions, however by applying a confidence interval and selecting from this range randomly when producing our Lambda max and SHI range estimates via stochastic simulations we account for this uncertainty in estimating adult survival in the absence of hunting mortality. Age at first reproduction (alpha) was also taken from Robinson (2005). Alternative sources of adult survival and age at reproduction estimates for birds are built into popharvest R package (Ernaud et al., 2021). This requires information on species specific body mass which we took from Robinson (2005). Where sex-specific body mass was reported we averaged male and female masses.

Short-term (2008-2018) and long-term (1970-2018) wintering population trends were taken from Burns et al. (2020), except for common snipe and Eurasian woodcock. Common snipe trends were taken from Woodward et al. (2020), with caution advised due to the small sample size. Resident woodcock long-term and short-term population trends were estimated at -29% for both on the basis of reported declines in breeding woodcock (Balmer et al., 2013; Heward et al., 2015). Migratory woodcock short-term and long-term trends were estimated at -11% and -22% on the basis of a 4-18% decline from 2008-2018 and an 11-33% decline from 1980-2018 (BirdLife International, 2021). Bird population status in the UK (Red/Amber/Green) was taken from Birds of Conservation Concern 5 (BoCC5; Stanbury et al. 2021).

**Table S1.** Assessment of realised values of recruitment, population growth or potential max population growth from demographic studies of brood success and survival for mallard and Canada goose across species range. These estimates are for hunted wild populations. Maximum population growth estimated by Demographic Invariant Method is 1.979 for mallard and 1.225 for Canada goose for comparison. “r max” is recruitment and where sex specific values given it is shown as Female|Male. If values are not sex dependent, single measure is reported. Age to which recruitment is assessed to in brackets, where 0+ is recruitment to autumn migration in ducks. Whether recruitment, population growth or both are estimated in the study they are highlighted in bold. Lambda max (λmax) is estimated from rmax by *exp*(rmax) where rmax is the only estimate in a given study. Productivity as number of fledged juveniles per female or per nest are not identical to recruitment (rmax) but have been provided to aid interpretation.

| **Species** | **Range** | **r max** | **λmax** | **Hunting mortality included** | **Reference** |
| --- | --- | --- | --- | --- | --- |
| Canada goose | NAmer\| resident | **0.31**(3)\| | 1.36 | Yes | Conover and Frank 2020 |
| Canada goose | NAmer \| all goose populations 1993-2019 average^a^ |  | **Mean: 1.074**  **Max: 1.24** | Yes | Sauer et al., 2020 |
| Canada goose | NAmer \| all goose populations 1968-2019 – annual rates^b,d^ |  | **Range of top 10: 1.92- 2.41**  **Median: 1.08; Mean: 1.10** | Yes | Sauer et al., 2020 |
| Canada goose | UK \|resident |  | **1.093** | Yes | Austin et al., 2007 |
| Canada goose | UK breeding index |  | **1.33-1.36** | Yes | Woodward et al. 2020 |
| Canada goose | UK wintering index |  | **1.58-1.74** | Yes | Woodward et al. 2020 |
| Canada goose | Demographic studies; %brood failure (e.g. 60%)*median clutch size (4-6) *%brood survival to autumn (0.40-0.86) | 1.38-1.8 |  | No | Ness et al., 2017; Cotter et al., 2014; Allan et al., 1995; Wright & Phillips, 1991; |
| Canada goose (England, Gravel pits) | Productivity (Mean autumn fledgings per nesting female)^e^ | 2.4-3.2;  3.18 (**1.2-1.6**) |  | No | Wright & Giles, 1988;  Wright & Phillips, 1991 |
| Canada goose | Young per adult 1970-74 (North Amer) | 0.29-1.18 |  |  | Raveling 1981 |
| Mallard | Finland\| wild | **0.21** (1yr from pulli) | 1.23 | Yes | Gunnarsson et al., 2008 |
| Mallard | NAmer\| migrants | **0.27** (0+) | 1.31 | No | Cowardin et al., 1985 |
| Mallard | UK breeding index |  | **1.23-1.33** | Yes | Woodward et al. 2020 |
| Mallard | UK wintering index |  | **1.18** (1.05 since 2000) | Yes | Woodward et al. 2020 |
| Mallard | NAmer \| all mallard populations 1968-2019 average^a^ |  | **Mean: 1.09**  **Max: 1.19** | Yes | Sauer et al., 2020 |
| Mallard | NAmer \| all Mallard populations 1968-2019 – annual rates^b,c^ |  | **Range of top 10: 1.3- 1.4**  **Median: 1.01; Mean: 1.01** |  | Sauer et al., 2020 |
| Mallard | NAmer\| maximum annual change in breeding population sizes 1955-2018 |  | **Regional Max: 1.59**  **Continental Max: 1.32** | Yes | Zimpfer et al., 2012 |
| Mallard | Demographic studies; %brood failure (e.g. 50%)*median clutch size (8-10) *%brood survival to autumn (0.23-0.44) | 1.58-2.1 |  |  | Danell and Sjöberg, 1979;  Petrie et al., 2000; Gunnarsson et al., 2008 |
| Mallard | Productivity (Mean autumn fledgings per nesting female)^e^ | 2.9;  2.2-2.7;  3.3  (**1.1-1.65**) |  |  | Petrie et al., 2000;  Majewski., 1986; Danell and Sjöberg, 1979. |

a) Average trend per year for net change between years stated.
b) Differential growth in annual index from NAmer Breeding bird survey for years stated.
c) Differential growth in annual change in index of abundance. Top 10 annual growth rates observed from seven NAmer states/territories between 1970 and 2014.
d) Differential growth in annual change in index of abundance. Top 10 annual growth rates observed from eight NAmer states/territories between 1982 and 2000.
e) Mean autumn fledgings per nesting female includes all failed nesting attempts and brood failures as well as duckling survival to independence but is not identical to rmax. Half of this productivity value, placed in bold, may be a suitable proxy of rmax but this is not recruitment to age at first reproduction.

Table S2: Estimated λ_max_, potential excess growth (PEG), sustainable hunt index (SHI) and probability of unsustainable harvest assuming “reported” adult survival and a “long” life history strategy

| **Species** | **Estimated λ_max_**  **(95% CI)** | **Potential Excess Growth**  **(95% CI)** | **Sustainable Hunt Index**  **(95% CI)** | **Probability of unsustainable harvest** |
| --- | --- | --- | --- | --- |
| Mallard | 1.841 (1.748 - 1.934) | 1,000,000 (920,000 – 1,100,000) | 0.957 (0.710 - 1.223) | 0.410 |
| Teal | 1.981 (1.761 - 2.204) | 150,000 (12,0000 – 170,000) | 0.992 (0.616 - 1.432) | 0.475 |
| Wigeon | 1.981 (1.814 - 2.149) | 150,000 (130,000 – 180,000) | 0.360 (0.146 - 0.610) | 0.000 |
| Gadwall | 1.675 (1.342 - 2.071) | 8000 (4500 – 11,000) | 0.820 (0.293 - 1.670) | 0.278 |
| Pintail | 1.774 (1.485 - 2.093) | 5700 (3800 - 7600) | 0.215 (0.043 - 0.443) | 0.000 |
| Shoveler | 1.909 (1.737 - 2.081) | 6300 (5200 - 7400) | 0.390 (0.150 - 0.664) | 0.000 |
| Tufted duck | 1.712 (1.647 - 1.781) | 38,000 (35,000 – 40,000) | 0.155 (0.076 - 0.237) | 0.000 |
| Pochard | 1.806 (1.722 - 1.888) | 1700 (1500 - 1900) | 0.334 (0.094 - 0.596) | 0.000 |
| Goldeneye | 1.603 (1.396 - 1.840) | 5000 (3500 - 6400) | 0.217 (0.029 - 0.456) | 0.000 |
| Canada goose | 1.252 (1.244 - 1.261) | 19,000 (17,000 – 20,000) | 1.829 (0.814 - 2.896) | 0.889 |
| Icelandic greylag goose | 1.203 (1.141 - 1.260) | 8400 (6000 – 11,000) | 2.738 (2.045 - 3.845) | 1.000 |
| British greylag goose | 1.203 (1.141 - 1.261) | 13,000 (9200 – 16,000) | 4.071 (0.853 - 8.099) | 0.952 |
| Pink-footed goose | 1.207 (1.189 - 1.225) | 48,000 (39,000 – 57,000) | 0.440 (0.071 - 0.861) | 0.000 |
| Resident woodcock | 1.424 (1.364 - 1.480) | 29,000 (20,000 – 39,000) | 0.672 (0.443 - 1.008) | 0.029 |
| Migratory woodcock | 1.424 (1.363 - 1.481) | 220,000 (150,000 – 300,000) | 0.604 (0.394 - 0.910) | 0.004 |
| Common snipe | 1.482 (1.385 - 1.562) | 220,000 (170,000 – 270,000) | 0.370 (0.137 - 0.659) | 0.000 |
| Golden plover | 1.676 (1.497 - 1.873) | 110,000 (80,000 – 130,000) | 0.021 (0.003 - 0.042) | 0.000 |

Table S3: Estimated λ_max_, potential excess growth (PEG), sustainable hunt index (SHI) and probability of unsustainable harvest assuming “reported” adult survival and a “short” life history strategy

| **Species** | **Estimated λ_max_**  **(95% CI)** | **Potential Excess Growth**  **(95% CI)** | **Sustainable Hunt Index**  **(95% CI)** | **Probability of unsustainable harvest** |
| --- | --- | --- | --- | --- |
| Mallard | 2.666 (2.436 - 2.914) | 1,600,000 (1,500,000 – 1800,000) | 0.596 (0.441 - 0.764) | 0.000 |
| Teal | 3.043 (2.479 - 3.686) | 240,000 (200,000 – 290,000) | 0.610 (0.374 - 0.895) | 0.003 |
| Wigeon | 3.044 (2.593 - 3.527) | 250,000 (210,000 – 300,000) | 0.222 (0.090 - 0.374) | 0.000 |
| Gadwall | 2.274 (1.555 - 3.313) | 13,000 (6800 – 19,000) | 0.522 (0.185 - 1.088) | 0.039 |
| Pintail | 2.501 (1.837 - 3.371) | 9200 (6000 – 13,000) | 0.136 (0.027 - 0.287) | 0.000 |
| Shoveler | 2.842 (2.417 - 3.322) | 10,000 (8300 – 12,000) | 0.241 (0.092 - 0.414) | 0.000 |
| Tufted duck | 2.346 (2.193 - 2.511) | 60,000 (55,000 – 64,000) | 0.098 (0.048 - 0.150) | 0.000 |
| Pochard | 2.576 (2.378 - 2.788) | 2700 (2400 - 3100) | 0.210 (0.059 - 0.372) | 0.000 |
| Goldeneye | 2.096 (1.661 - 2.664) | 7800 (5300 – 10,000) | 0.140 (0.019 - 0.296) | 0.000 |
| Canada goose | 1.294 (1.284 - 1.305) | 21,000 (20,000 – 23,000) | 1.600 (0.706 - 2.525) | 0.818 |
| Icelandic greylag goose | 1.233 (1.159 - 1.304) | 9500 (6700 – 12,000) | 2.417 (1.783 - 3.403) | 1.000 |
| British greylag goose | 1.234 (1.161 - 1.305) | 15,000 (10,000 – 19,000) | 3.587 (0.742 - 7.250) | 0.933 |
| Pink-footed goose | 1.238 (1.216 - 1.259) | 54,000 (44,000 – 65,000) | 0.386 (0.062 - 0.761) | 0.000 |
| Resident woodcock | 1.553 (1.462 - 1.639) | 36,000 (24,000 – 49,000) | 0.540 (0.354 - 0.815) | 0.000 |
| Migratory woodcock | 1.553 (1.467 - 1.638) | 270,000 (180,000 – 370,000) | 0.483 (0.315 - 0.724) | 0.000 |
| Common snipe | 1.641 (1.493 - 1.771) | 270,000 (210,000 – 340,000) | 0.293 (0.109 - 0.525) | 0.000 |
| Golden plover | 2.258 (1.873 - 2.735) | 170,000 (120,000 – 210,000) | 0.013 (0.002 - 0.027) | 0.000 |

Table S4: Estimated λ_max_, potential excess growth (PEG), sustainable hunt index (SHI) and probability of unsustainable harvest assuming “estimated” adult survival and a “long” life history strategy

| **Species** | **Estimated λ_max_** | **Potential Excess Growth**  **(95% CI)** | **Sustainable Hunt Index**  **(95% CI)** | **Probability of unsustainable harvest** |
| --- | --- | --- | --- | --- |
| Mallard | 1.435 | 590,000 (590,000 – 600,000) | 1.619 (1.221 - 2.006) | 1.000 |
| Teal | 1.515 | 90,000 (87,000 – 94,000) | 1.623 (1.062 - 2.197) | 1.000 |
| Wigeon | 1.461 | 85,000 (76,000 – 95,000) | 0.652 (0.267 - 1.068) | 0.072 |
| Gadwall | 1.458 | 5800 (5700 - 6000) | 1.052 (0.441 - 1.677) | 0.535 |
| Pintail | 1.454 | 3700 (3300 - 4200) | 0.322 (0.067 - 0.601) | 0.000 |
| Shoveler | 1.470 | 3800 (3400 - 4100) | 0.653 (0.253 - 1.078) | 0.081 |
| Tufted duck | 1.458 | 26,000 (26,000 – 27,000) | 0.222 (0.108 - 0.335) | 0.000 |
| Pochard | 1.442 | 1100 (960 - 1200) | 0.540 (0.151 - 0.948) | 0.005 |
| Goldeneye | 1.449 | 3900 (3700 - 4100) | 0.271 (0.038 - 0.506) | 0.000 |
| Canada goose | 1.164 | 12,000 (12,000 – 13,000) | 2.724 (1.198 - 4.278) | 1.000 |
| Icelandic greylag goose | 1.170 | 7100 (6900 - 7400) | 3.156 (2.788 - 3.547) | 1.000 |
| British greylag goose | 1.170 | 11,000 (11,000 – 11,000) | 4.794 (1.022 - 8.484) | 0.978 |
| Pink-footed goose | 1.175 | 41,000 (34,000 – 48,000) | 0.512 (0.082 - 0.999) | 0.025 |
| Resident woodcock | 1.302 | 22,000 (15,000 – 28,000) | 0.896 (0.600 - 1.322) | 0.286 |
| Migratory woodcock | 1.302 | 160,000 (110,000 – 210,000) | 0.802 (0.535 - 1.185) | 0.150 |
| Common snipe | 1.336 | 160,000 (130,000 – 180,000) | 0.495 (0.188 - 0.856) | 0.000 |
| Golden plover | 1.545 | 89,000 (80,000 – 98,000) | 0.024 (0.003 - 0.046) | 0.000 |

Table S5: Estimated λ_max_, potential excess growth (PEG), sustainable hunt index (SHI) and probability of unsustainable harvest assuming “estimated” adult survival and a “short” life history strategy

| **Species** | **Estimated λ_max_** | **Potential Excess Growth**  **(95% CI)** | **Sustainable Hunt Index**  **(95% CI)** | **Probability of unsustainable harvest** |
| --- | --- | --- | --- | --- |
| Mallard | 1.738 | 910,000 (900,000 – 910,000) | 1.057 (0.799 - 1.311) | 0.605 |
| Teal | 1.901 | 140,000 (130,000 – 150,000) | 1.051 (0.683 - 1.426) | 0.571 |
| Wigeon | 1.791 | 130,000 (120,000 – 150,000) | 0.422 (0.171 - 0.693) | 0.000 |
| Gadwall | 1.784 | 9000 (8700 – 9200) | 0.693 (0.288 - 1.095) | 0.140 |
| Pintail | 1.775 | 5700 (5100 - 6400) | 0.211 (0.044 - 0.388) | 0.000 |
| Shoveler | 1.809 | 5800 (5200 - 6300) | 0.420 (0.161 - 0.702) | 0.000 |
| Tufted duck | 1.784 | 41,000 (40,000 – 41,000) | 0.144 (0.070 - 0.218) | 0.000 |
| Pochard | 1.752 | 1600 (1500 - 1800) | 0.351 (0.098 - 0.621) | 0.000 |
| Goldeneye | 1.766 | 6000 (5600 - 6300) | 0.176 (0.026 - 0.330) | 0.000 |
| Canada goose | 1.186 | 14,000 (13,000 – 15,000) | 2.437 (1.075 - 3.810) | 0.995 |
| Icelandic greylag goose | 1.193 | 8000 (7800 - 8300) | 2.796 (2.466 - 3.142) | 1.000 |
| British greylag goose | 1.193 | 12,000 (12,000 – 13,000) | 4.200 (0.888 - 7.518) | 0.960 |
| Pink-footed goose | 1.200 | 46,000 (39,000 – 54,000) | 0.456 (0.075 - 0.881) | 0.000 |
| Resident woodcock | 1.379 | 27,000 (19,000 – 35,000) | 0.737 (0.495 - 1.084) | 0.074 |
| Migratory woodcock | 1.379 | 200,000 (140,000 – 260,000) | 0.656 (0.440 - 0.965) | 0.013 |
| Common snipe | 1.427 | 200,000 (170,000 – 230,000) | 0.411 (0.153 - 0.706) | 0.000 |
| Golden plover | 1.966 | 140,000 (120,000 – 150,000) | 0.015 (0.002 - 0.030) | 0.000 |
